# Partial prevention of glucocorticoid-induced osteocyte deterioration with osteocrin gene therapy

**DOI:** 10.1101/2021.11.19.469265

**Authors:** Courtney M. Mazur, Christian D. Castro Andrade, Tadatoshi Sato, Michael Bruce, Mary L. Bouxsein, Jialiang S. Wang, Marc N. Wein

**Affiliations:** Endocrine Unit, Massachusetts General Hospital, Harvard Medical School, Boston, MA; Center for Advanced Orthopedic Studies, Department of Orthopedic Surgery, Beth Israel Deaconess Medical Center, Harvard Medical School, Boston, Massachusetts, USA; Broad Institute of Harvard and MIT, Cambridge, MA; Harvard Stem Cell Institute, Cambridge, MA

## Abstract

Glucocorticoid (GC)-induced osteoporosis and subsequent bone fragility are preceded by death and dysfunction at the cellular level. In particular, short-term glucocorticoid excess suppresses osteocyte remodeling of the surrounding bone mineral, causes apoptosis of osteoblasts and osteocytes, and disrupts homeostatic bone remodeling. Preventing apoptosis and preserving osteocyte morphology and function could be effective means of preventing bone loss during glucocorticoid excess. We hypothesized that osteocrin, which preserves osteocyte viability and morphology in other models where osteocyte defects exist, could prevent osteocyte death and dysfunction in a GC excess model. We used a liver-targeted adeno-associated virus (AAV8) to induce osteocrin overexpression in mice one week prior to implantation with prednisolone or placebo pellets. After 28 days, tissues were collected for micro-CT and histological analysis. GC excess caused the expected reduction in cortical bone thickness and osteocyte canalicular length in control AAV8-treated mice, and these effects were blunted in mice overexpressing osteocrin. However, GC-induced changes in cortical porosity, trabecular bone mass, and gene expression were not prevented by osteocrin. While the mechanism of osteocrin’s effects on osteocyte morphology warrants further investigation, this study does not support a role for this model of osteocrin supplementation to combat the full skeletal effects of GC excess.

## Introduction

Long-term glucocorticoid (GC) treatment is a well-known cause of drug-induced osteoporosis and osteonecrosis. By reducing bone formation, increasing bone resorption, and reducing bone quality, glucocorticoids increase the risk of fragility fractures in a dose and duration-dependent manner (1). Due to the usefulness of glucocorticoids in treating patients with a wide variety of inflammatory conditions, many studies have investigated the effects of antiresorptives and anabolic therapies for preserving bone mass during GC use. Bisphosphonates, denosumab, and teriparatide show some efficacy in improving bone mass and reducing fracture risk in patients receiving GCs (2–6). However, these treatments are accompanied by rare but serious side effects, such as increased risk of osteonecrosis, atypical fractures, and hypercalcemia (5,7,8), so improved therapies are needed. Furthermore, available therapies for GC-induced osteoporosis largely ignore osteocytes.

Healthy osteocytes play key roles in bone by secreting paracrine factors to coordinate the activity of osteoblasts and osteoclasts and by remodeling the bone matrix surrounding their lacuno-canalicular network. This local remodeling maintains organization of the lacuno-canalicular network and contributes to bone material quality; genetic and pharmacological interventions that suppress osteocyte remodeling activity reduce bone quality and increase fragility (9–11). A large body of research demonstrates that high-dose glucocorticoid treatment induces osteocyte death in humans and in mice (12–17). Even before causing apoptosis, GC excess reduces the number and length of osteocyte dendritic projections and disrupts the ability of osteocytes to remodel the lacuno-canalicular network (14,18,19). Preventing osteocyte apoptosis, preserving their dendritic phenotype, and maintaining remodeling activity during GC excess could be a novel method of combatting GC-induced osteoporosis and fragility.

Recent work in our lab revealed a role for the transcription factor Sp7 in osteocyte development (20). Conditional ablation of Sp7 causes reduced osteocyte dendrite number and length, apoptosis of embedded osteocytes, and increased cortical porosity and low bone mass in young mice. This phenotype can be rescued by overexpression of one of the genes whose levels are reduced by Sp7 deletion – osteocrin. Osteocrin (Ostn) is a secreted peptide that occupies the clearance receptor NPR-C on target cells, thereby preventing clearance of CNP and allowing increased CNP signaling through NPR-B (21). Downstream of NPR-B, CNP signaling leads to production of cGMP, activation of protein kinase G, and action on a variety of targets including Erk phosphorylation and modulation of the cytoskeleton through RhoA and myosin light chain phosphatase (22). Since Ostn over-expression was able to rescue osteocyte morphology and survival defects in Sp7 mutant mice, we hypothesized that overexpression of Ostn could prevent the changes in osteocyte morphology and viability in response to GCs.

Here we tested the influence of systemically-administered Ostn via AAV8 gene therapy on the skeletal effects of GCs in mice. We report that AAV8-Ostn treatment blunted GC-induced effects such as reduction in cortical bone mass and osteocyte canalicular length. Despite these apparently beneficial findings we noted overall mild effects of GC treatment in this model, and AAV8-Ostn therapy failed to rescue other skeletal consequences of GCs such as increased cortical bone porosity and reduced trabecular bone mass._Taken together, these data demonstrate potential beneficial effects of AAV8-Ostn therapy for osteocytes that may not fully combat this common cause of skeletal fragility._

## Methods

### Animals

6-week-old male C57BL/6J and FVB mice were purchased from Jackson Labs (#000664 and #001800). Mice were randomized into four weight-matched groups at 7 weeks old and injected intraperitoneally with either AAV8-CAG-mOstn-WPRE or AAV8-CAG-eGFP in 100 uL saline (5*10^11^ genome copies/mouse, Vector Biolabs). For pilot experiments in C57BL/6J mice, AAV8-Ostn-treated mice were euthanized at 24 hours, 3 days, and 7 days-post-injection for quantification of *Ostn* expression in liver.

8 week old FVB mice treated with each AAV were then implanted with slow-release prednisolone or placebo pellets giving an average dose of 2.8 mg/kg/day (4.3 mg; 60-day release; Innovative Research of America). Before the procedure, mice were anesthetized with isofluorane and given one dose of long-acting buprenorphine (0.08 mg/kg). Fur was shaved, and the area was prepared with isopropanol and betadine. Pellets were implanted subcutaneously at the base of the neck using a trochar and the incision sealed with cyanoacrylate tissue glue. Mice were observed daily for three days following the procedure and then 3-4 times per week. One mouse (prednisolone+AAV8-Ostn group) was euthanized early due to wound dehiscence and removed from the study. Four weeks after pellet implantation, mice were euthanized with CO_2_ followed by cervical dislocation, and tissues were collected for analysis. All procedures involving animals were performed in accordance with guidelines issued by the Institutional Animal Care and Use Committees (IACUC) in the Center for Comparative Medicine at the Massachusetts General Hospital and Harvard Medical School under approved Animal Use Protocols (2019N000201). All animals were housed in the Center for Comparative Medicine at the Massachusetts General Hospital (21.9 ± 0.8 °C, 45 ± 15% humidity, and 12-h light cycle 7 am–7 pm).

### Serum

Just prior to euthanasia, whole blood was collected via cheek bleed into serum separator tubes (BD 365967). Blood was allowed to clot for at least 10 minutes, then centrifuged for 10 minutes at 4 °C to separate the serum. Serum was stored at −80°C until analysis by procollagen type 1 N-terminal peptide (P1NP) and C-terminal collagen crosslinks (CTX) assays (IDS #AC-33F1 and #AC-06F1) based on the manufacturer’s instructions.

### RNA

Liver and bone were collected for RNA immediately following euthanasia. A portion of the liver was flash-frozen in liquid nitrogen and stored at −80 °C. Liver tissue was pulverized in TRIzol and RNA isolated using chloroform followed by precipitation in isopropanol. Humeri were dissected and cleaned of muscle. The epiphyses were trimmed, bones were transferred into microcentrifuge tubes, and marrow was removed by centrifugation (17,000G for 15s at RT). Bones were then snap-frozen in liquid nitrogen and stored in liquid nitrogen. Bones were pulverized in TRIzol and RNA extracted using chloroform followed by precipitation in isopropanol and column purification (PureLink RNA Mini Kit, Invitrogen). Two humeri (placebo+AAV8-control and prednisolone+AAV8-control) were mislabeled and excluded from further analysis. RNA was DNase treated and reverse transcribed to cDNA (PrimeScript, Takara) before RT-qPCR with SYBR primers (Table 1) on an Applied Biosystems StepOnePlus system.

**Table 1:**
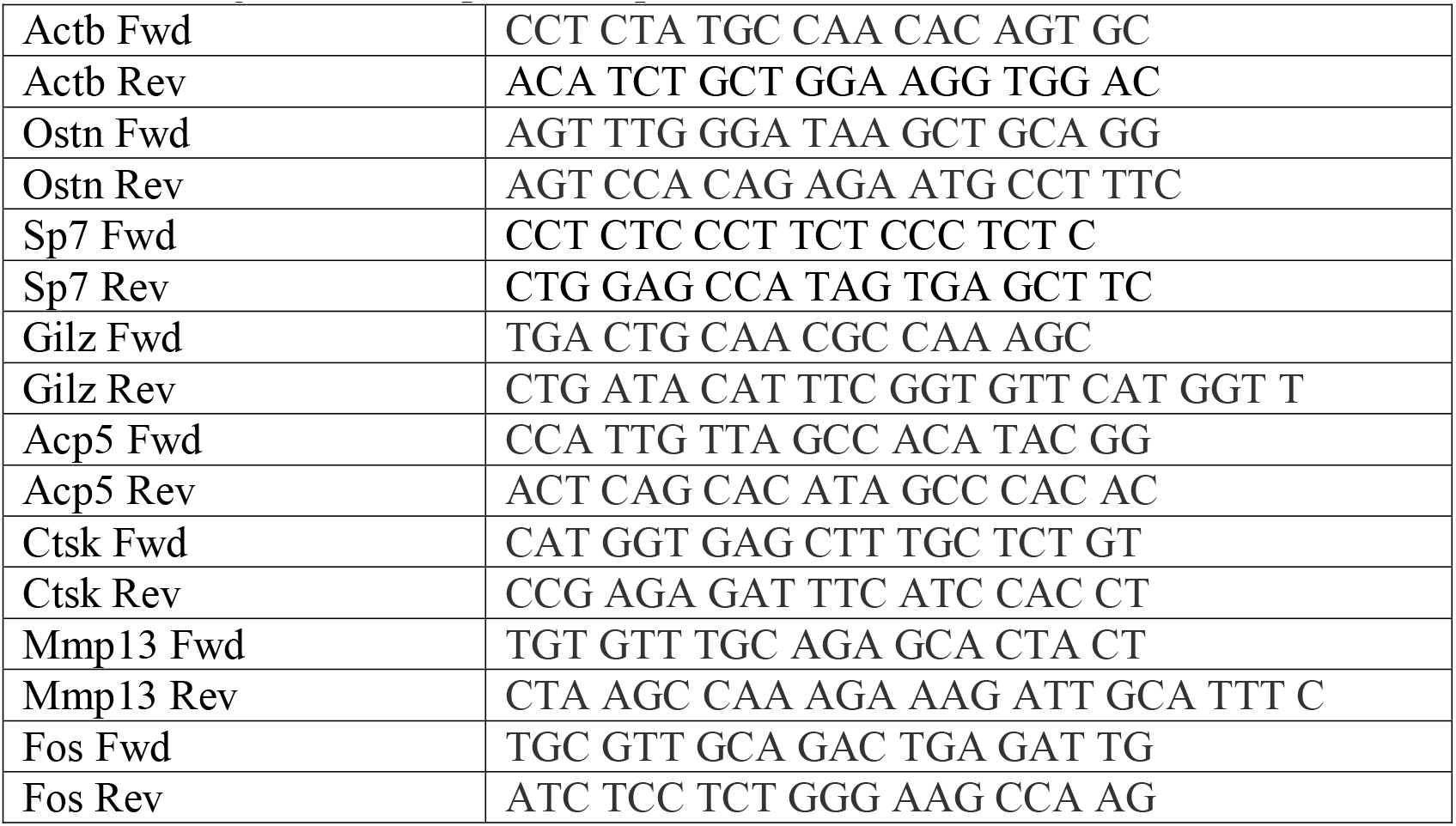
Oligonucleotide primer sequences

### Micro-CT

Femurs were dissected and stored in cold 70% ethanol until scanning. Micro-CT imaging was performed on a bench-top scanner (μCT40, Scanco Medical AG, Brüttisellen, Switzerland) to measure the morphology of the femoral mid-diaphysis and distal metaphysis. A 500 μm region of the femoral mid-diaphysis was scanned using 70 kVp peak X-ray tube potential, 113 mA X-ray tube current, 200 ms integration time, and 10 μm isotropic voxels. Cortical bone was segmented from soft tissue with a threshold of 700 mg HA/cm^3^. A 1500 μm region of the femoral mid-diaphysis, starting 1700 μm proximal to the distal growth place and extending distally, was scanned using the same settings. Cancellous bone was segmented from soft tissue using a threshold of 300 mg HA/cm^3^ and separated from cortical bone with a semiautomated contouring program. Scanning and analysis was performed in the Scanco Evaluation Program in a blinded manner and adhered to the guidelines for the use of micro-CT for the assessment of bone architecture in rodents (23).

### Histology

Tibias were dissected and fixed in 10% neutral-buffered formalin for 24 hours. Bones were decalcified in EDTA for two weeks, then one bone was dehydrated in graded ethanols and embedded in paraffin. The contralateral tibia was incubated in 30% sucrose solution and embedded in OCT. 5 μm longitudinal sections were obtained from paraffin blocks and 10 μm sections were obtained from frozen blocks.

Frozen sections were washed in PBS, post-fixed in 4% paraformaldehyde (PFA), permeabilized in 0.05% saponin, and stained with phalloidin for 30 minutes (1:1000, Abcam ab112125). After washing, samples were incubated with DAPI (Invitrogen, 5 μg/mL) for 15 minutes, washed again, and coverslips mounted with Fluoromount-G. Imaging was conducted on a Zeiss Airyscan confocal microscope with a 40x oil-immersion objective.

Paraffin sections were deparaffinized in three changes of xylene and hydrated through graded ethanols, then used for hematoxylin & eosin (H&E), terminal deoxynucleotidyl transferase dUTP nick end labeling (TUNEL), RNAscope, silver nitrate, and tartrate-resistant acid phosphatase (TRAP). H&E staining was performed using standard protocols, followed by imaging on a Keyence BZ-X710 microscope at 20x.

For TUNEL, rehydrated sections were post-fixed in 4% PFA and permeabilized with PCR-grade Proteinase K (20 μg/mL) for 30 minutes at 37 °C. Samples were incubated with TUNEL reaction mixture (Roche 11684795910) for 60 minutes at 37 °C followed by DAPI, then mounted with Fluoromount-G. Fluorescent images were captured at 20x on a Nikon Eclipse Ni microscope.

RNA in situ hybridization for Ostn was performed with RNAscope 2.5 HD Assay-Brown (Advanced Cell Diagnostics). Tibia sections were prepared by deparaffinization in fresh xylene and 100% ethanol, then dried in a 60 °C oven for 30 minutes. Samples were incubated with hydrogen peroxide at room temperature for 10 minutes, then with pepsin for 30 minutes at 40 °C (Sigma R2283). After washing, samples were incubated with target probe (Ostn, NM_198112.2, region 2-1144) and amplified following the manufacturer’s instructions. Signal was detected using DAB for 10 minutes, then sections were briefly counterstained with 50% hematoxylin, dehydrated, cleared, and mounted. Brightfield images were captured at 20x on a Nikon Eclipse Ni microscope.

Silver nitrate staining solution was freshly prepared from two parts 50% silver nitrate and one part 2% gelatin type A with 1% formic acid (18). Sections were incubated for 55 minutes in the dark, then washed and incubated in 5% sodium thiosulfate for 10 minutes. After dehydrating and mounting, images were captured on a Keyence microscope system at 40x.

For TRAP staining, sections were rehydrated to water, then incubated in 0.2 M acetate buffer (pH 5.0) for 30 minutes. TRAP staining solution was freshly prepared with 0.5 mg/mL Napthol As-MX Phosphate and 1.1 mg/mL Fast Red TR salt in 0.2 M acetate buffer, and samples were incubated for 30 minutes at 60 °C. Slides were then rinsed in water, counterstained in Fast Green for 1 minute, dipped in 1% acetic acid, rinsed in water, and mounted with aqueous medium. Imaging was conducted on a Nanozoomer slide scanner (Hamamatsu).

### Image quantification

All analyses were performed on a blinded basis using images from uniform areas from at least four regions of interest (ROIs) representing the anterior and posterior cortex of the tibia. The number of osteocyte nuclei and empty lacunae was counted in four H&E stained images per bone to calculate the proportion of empty lacunae. For TUNEL and TRAP staining, the proportion of positively-stained osteocytes was calculated relative to total DAPI-stained nuclei and total cortical bone area, respectively.

In silver nitrate-stained bones, the length of ten dendrites on each of three osteocytes in all four ROIs was measured using ImageJ (120 dendrites/mouse). Additionally, the total number of dendrites was counted for five representative osteocytes in all four ROIs. For phalloidin-stained bones, signal from osteocyte cell bodies was removed using DAPI as a mask, and images were uniformly thresholded to remove background. The density of osteocyte dendrites was calculated as the percent of image area stained by phalloidin.

### Cell culture

Ocy454 cells were maintained in alpha-minimum essential medium (MEM) supplemented with heat-inactivated 10% fetal bovine serum and 1% antibiotic–antimycotic (Gibco) at 33 °C with 5% CO_2_. Cells are transferred to 37 °C to inactivate the temperature-sensitive T antigen and promote osteocytic differentiation. They were routinely assessed by Sost and Dmp1 expression at 37 °C as previously described (24,25).

For gene expression analysis, cells were grown in 24-well plates at 33 °C until confluent, then differentiated at 37 °C for 7 days. Dexamethasone (Sigma D2915) was prepared in water and added to culture media at the indicated final concentrations, with water vehicle added to control groups at equal volume. After three hours of treatment cells were washed and lysed in RLT. Lysate was passed through QIAshredder columns (Qiagen), then RNA was purified with PureLink columns (Invitrogen) and reverse transcribed to cDNA as above.

### Statistics

For analysis of *in vivo* gene expression data (2 groups), groups were compared using unpaired 2-tailed t-tests. For *in vitro* gene expression data (3 groups), analysis was performed using 1-way ANOVA followed by pairwise comparisons of groups with Tukey’s multiple comparisons tests. To analyze the effects of prednisolone and osteocrin on bone, two-way ANOVA was performed, followed by Sidak’s multiple comparisons tests between placebo- and prednisolone-treated mice within the same AAV8 treatment group. The ROUT method (Q=1%) was used to remove outliers from each analysis (GraphPad Prism 8.4). P values less than 0.05 were considered to be significant. Each plotted data point represents one independent biological replicate.

## Results

The goal of this study was to assess the therapeutic potential of osteocrin (Ostn) overexpression on GC-induced skeletal disease. To establish a rationale for this study, we first tested the effects of GCs on Sp7 and Ostn expression in an osteocyte cell culture model. Dexamethasone treatment of Ocy454 cells reduced both *Sp7* and *Ostn* mRNA expression in a dose dependent manner while causing the expected increase in the canonical glucocorticoid receptor target gene *Gilz* (26) (Figure 1A-C).

**Figure 1:**
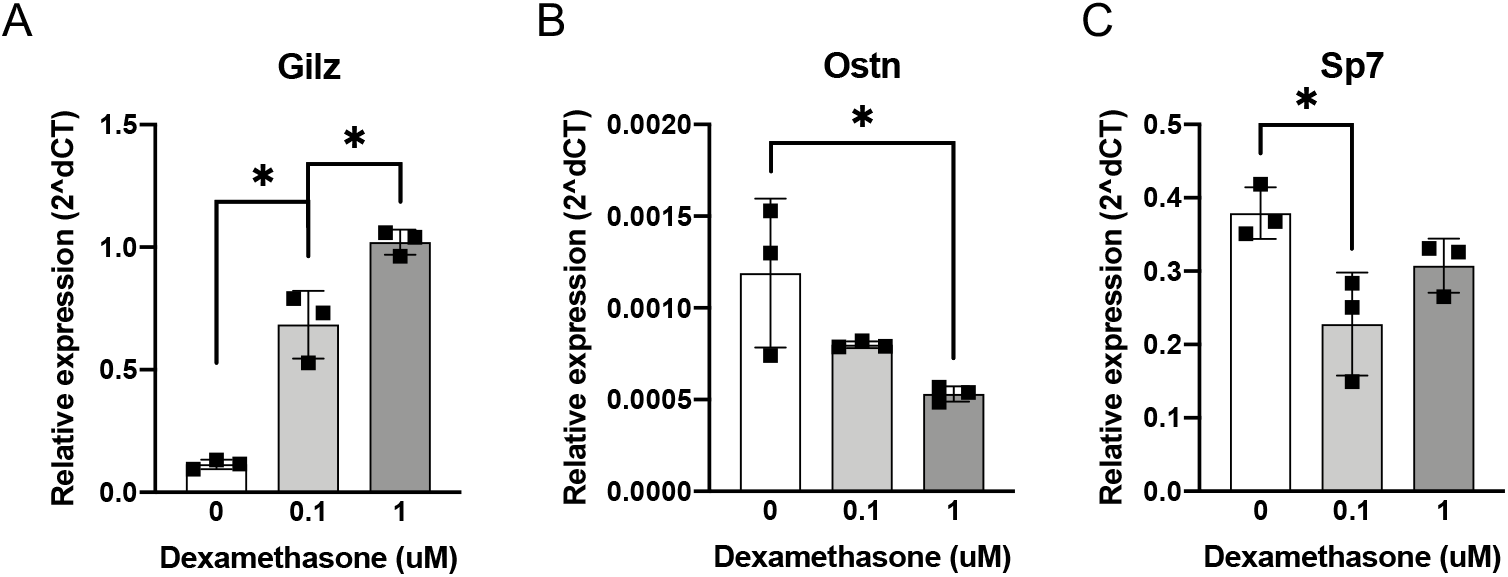
Glucocorticoids suppress *in vitro* Ostn expression. Ocy454 cells were treated with dexamethasone at the indicated concentrations for 3 hours. Expression of (A) canonical glucocorticoid receptor target gene *Gilz*, (B) *Ostn*, and (C) transcription factor *Sp7* were measured by RT-qPCR and presented relative to *Actb*. n=3. *p<0.05. Error bars show +/- SD.

To determine the short-term kinetics of *Ostn* expression *in vivo* in response to administration of the AAV8-Ostn vector, we first conducted a pilot experiment in C57BL/6J mice. Importantly, AAV8 has strong tropism for liver; therefore, this system delivers Ostn cDNA for hepatic expression with subsequent delivery of osteocrin protein to bone via the circulation (27,28). Mice were injected intraperitoneally with AAV8-Ostn and livers collected 1, 3, or 7 days later. Expression of hepatic *Ostn* mRNA increased rapidly over the first three days of AAV8 infection and remained high at 7 days (Supp Fig 1A). To allow for protein translation and transport to bone via the circulation, a seven-day lead time was chosen for AAV8-Ostn injection before starting glucocorticoid treatment.

One week after injection with AAV8-Ostn or AAV8-control vector, FVB mice (a strain reported to show robust glucocorticoid-induced bone loss (18,29,30) were implanted subcutaneously with continuous-release placebo or prednisolone pellets (2.8 mg/kg/d). Weekly monitoring of body weight showed that prednisolone-treated mice initially lost weight and maintained average body weights below those of placebo-treated mice (Supp Fig 1C-D). Weight loss due to muscle atrophy is a known side effect of high-dose glucocorticoid treatment in mice (31), so these findings suggest efficacy of the glucocorticoid excess model. Five weeks after infection, hepatic *Ostn* expression remained elevated for mice implanted with both placebo and prednisolone pellets (Supp Fig 1B), consistent with the documented persistence of AAV-mediated gene expression (27,28).

Similar to *in vitro* dexamethasone treatments, prednisolone reduced *Ostn* mRNA in humeri of AAV8-control-treated mice, though at this time point there was no detectable effect on *Sp7* mRNA in whole cortical bone samples (Fig 2A). The suppression of Ostn by prednisolone *in vivo* further supports the rationale for our hypothesis that Ostn overexpression could rescue the deleterious skeletal effects of glucocorticoids. In contrast to previous mRNA analyses after 7 days of prednisolone treatment (18), we did not find induction of glucocorticoid-responsive *Gilz* or suppression of proteinase *Mmp13* 28 days after prednisolone pellet implantation here. However, we did detect significant induction of osteoclast marker genes *Acp5* (encodes TRAP5b) and *Ctsk* (encodes Cathepsin K) by prednisolone (Fig 2A). Prednisolone-induced increases in *Ctsk* and *Acp5* were not prevented by AAV8-Ostn treatment (Fig 2B).

**Figure 2:**
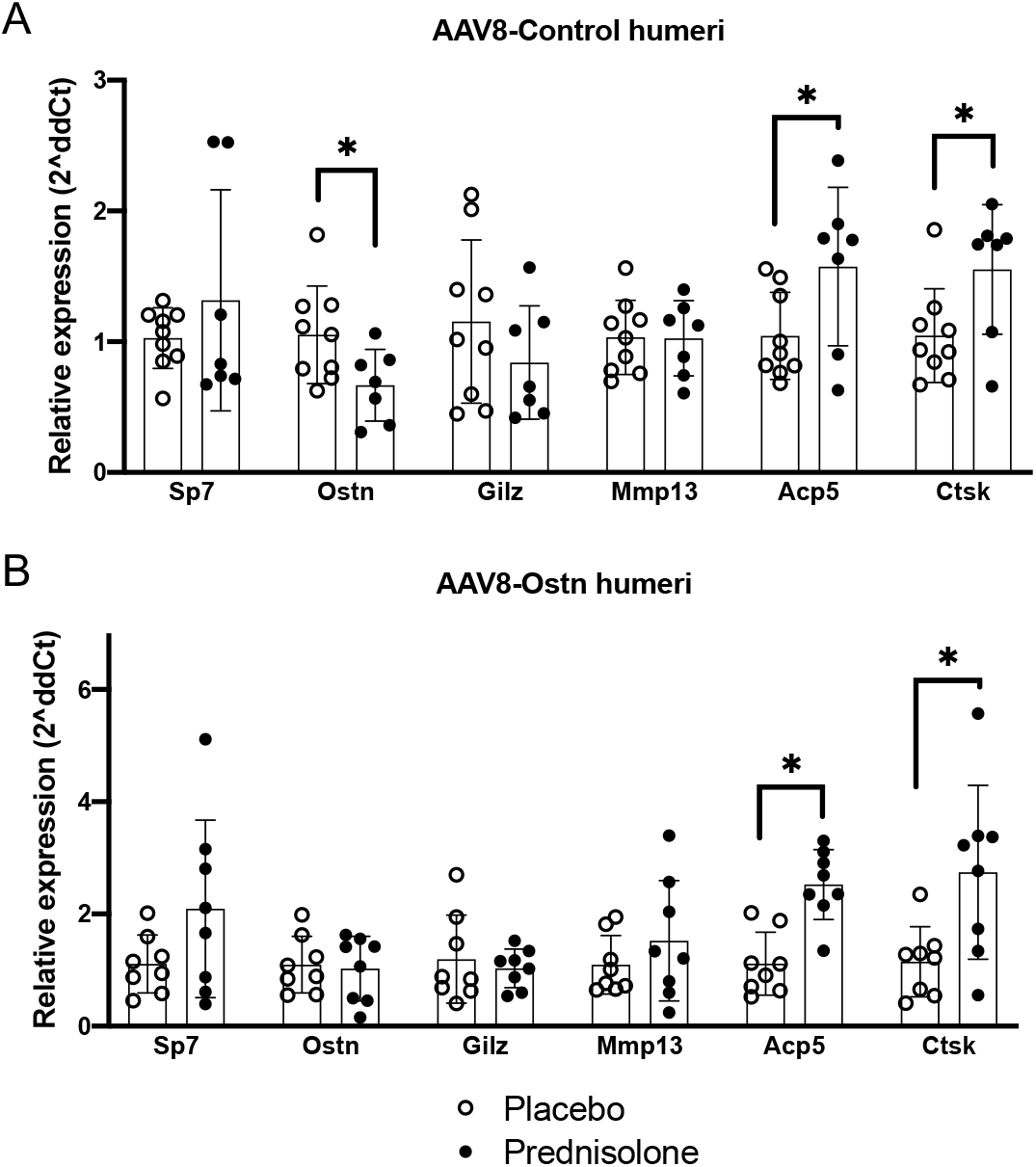
Prednisolone-induced changes in bone gene expression. Gene expression was measured relative to *Actb* in humeri after 28 days of prednisolone or placebo treatment in (A) AAV8-Control and (B) AAV8-Ostn-treated mice. *p<0.05 compared to corresponding placebo group. Error bars show +/- SD.

We previously reported that Ostn peptide enhances Erk1/2 phosphorylation in cultured osteoblasts and osteocytes (20), and here we show modest induction of the Erk1/2-responsive gene *Fos* (encodes cFos) by AAV8-Ostn treatment in mice (Supp Fig 2A). Bone *Fos* expression is significantly correlated with liver *Ostn* expression (Supp Fig 2B), suggesting that the Ostn peptide encoded by AAV8-Ostn administration induces signaling and downstream gene expression changes in bone. Further supporting the efficacy of AAV8-Ostn treatment, comparison of all AAV8-control to all AAV8-Ostn-treated mice reveals a significant reduction in femoral cortical porosity with AAV8-Ostn (Fig 3C, main effect of two-way ANOVA analysis), which is consistent with our previous studies in Sp7-deficient mice (20).

**Figure 3:**
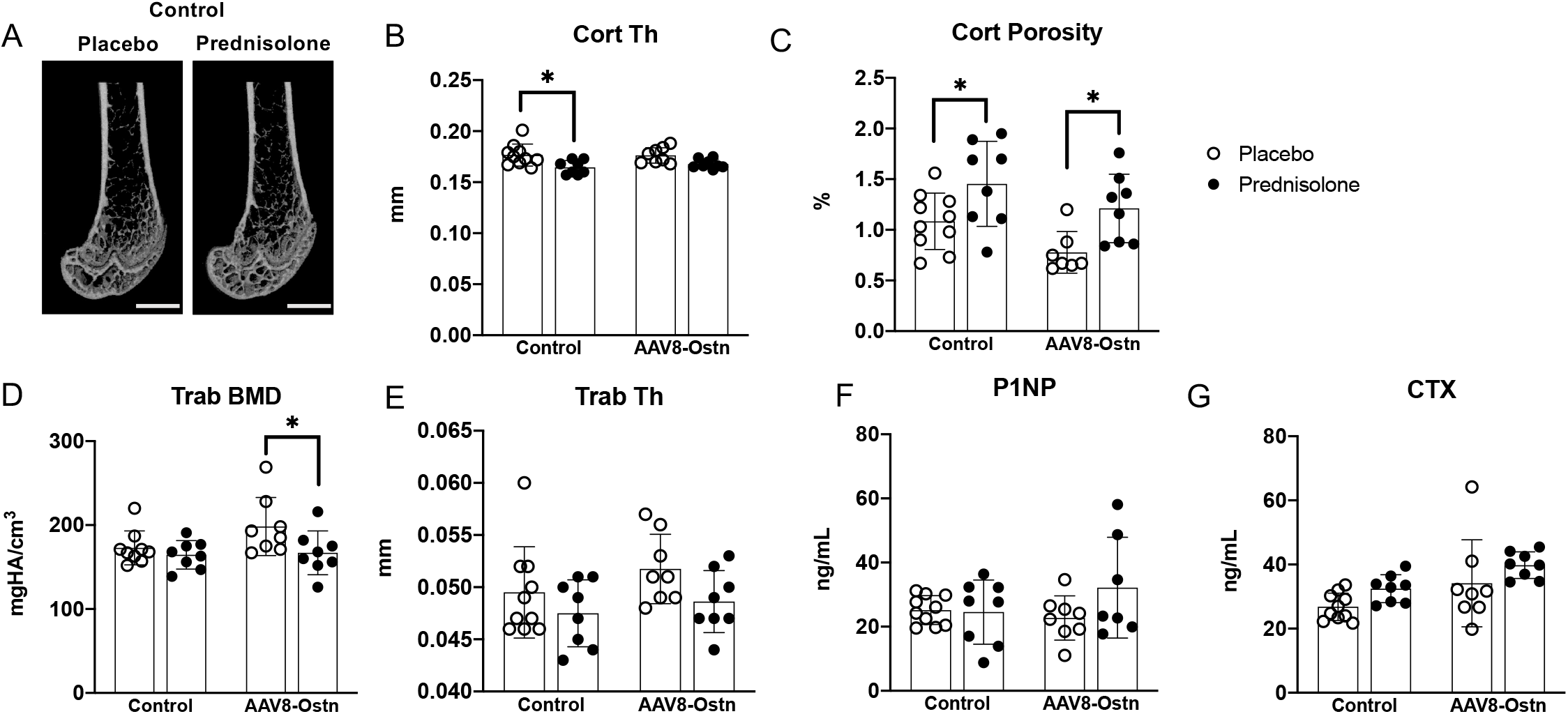
Prednisolone exerts a mild overall effect on bone mass, formation, and resorption. (A) Representative micro-CT images of the distal femur in AAV8-Control-treated mice after 28 days of placebo or prednisolone treatment. Scale bar = 1 mm. (B) Cortical thickness (Cort Th) and (C) Cortical porosity (Cort porosity, one outlier removed from placebo+AAV8-Ostn group) were measured at the femoral mid-diaphysis. (D) Trabecular bone mineral density (Trab BMD, one outlier removed from placebo+AAV8-Control group) and (E) Trabecular thickness (Trab Th) were measured in the distal femoral metaphysis. (F) Serum biomarker of bone formation, procollagen type 1 N-terminal peptide (P1NP, one outlier removed from prednisolone+AAV8-Ostn group). (G) Serum biomarker of bone formation, C-terminal collagen crosslinks (CTX). *p<0.05 compared to corresponding placebo group. All error bars show +/- SD.

In the femur, prednisolone caused the expected reduction in mid-diaphyseal cortical thickness in AAV8-control-treated mice (18,32). In contrast, the effect of prednisolone on cortical thickness did not reach statistical significance in AAV8-Ostn-treated mice (Fig 3A, B). However, prednisolone significantly increased cortical porosity and decreased cortical area fraction in both AAV8-control and AAV8-Ostn-treated mice (Fig 3C, Table 2). We also found that prednisolone reduced trabecular thickness, reduced trabecular bone mineral density (BMD), and increased trabecular structure model index (SMI), but these effects were not mitigated by AAV8-Ostn (Fig 3D, E, Table 2).

**Table 2:** Microstructural properties of the femoral mid-diaphysis and metaphysis. Properties measured in the femoral metaphysis include bone volume fraction (BV/TV), bone mineral density (BMD), trabecular number (Tb. N.), trabecular thickness (Tb. Th.), trabecular separation (Tb. Sp.) and structural model index (SMI). Properties measured at the femoral mid-diaphysis include cortical thickness (Ct. Th.), bone area fraction (Ct. At./Tt. Ar.), total mineral density (Ct. TMD), and cortical porosity (Ct. Porosity). All values presented as mean +/-SD. ^ One outlier removed. $ Overall effect of prednisolone p<0.05, Two-way ANOVA. + Overall effect of osteocrin p<0.05, Two-way ANOVA. * p<0.05 compared to corresponding placebo group.

|  | Placebo+Control<br>(n=10) | Placebo+<br>AAV8-Ostn (n=8) | Prednisolone+Control<br>(n=8) | Prednisolone+<br>AAV8-Ostn (n=8) |
| --- | --- | --- | --- | --- |
| Bone length<br>(mm) | 14.56 +/- 0.26 | 14.64 +/- 0.46 | 14.35 +/- 0.30 | 14.37 +/- 0.57 |
| BV/TV (%) | 16.44 +/- 2.67 ^ | 20.61 +/- 5.38 | 15.56 +/- 2.55 | 16.65 +/- 3.80 |
| BMD<br>(mgHA/cm<sup>3</sup>) | 173 +/- 20 ^ | 198 +/- 35 | 164 +/- 17 \$ | 167 +/- 26 \$* |
| Tb. N. (1/mm) | 5.203 +/- 0.514 | 5.479 +/- 0.618 | 5.036 +/- 0.575 | 5.119 +/- 0.685 |
| Tb. Th. (mm) | 0.049 +/- 0.004 | 0.052 +/- 0.003 | 0.047 +/- 0.003 \$ | 0.049 +/- 0.003 \$ |
| Tb. Sp. (mm) | 0.192 +/- 0.020 | 0.183 +/- 0.021 | 0.200 +/- 0.023 | 0.196 +/- 0.030 |
| SMI | 1.951 +/- 0.195 ^ | 1.621 +/- 0.494 | 2.106 +/- 0.253 \$ | 2.021 +/- 0.335 \$* |
| Ct. Th. (mm) | 0.177 +/- 0.011 | 0.176 +/- 0.008 | 0.164 +/- 0.007 \$* | 0.168 +/- 0.005 \$ |
| Ct. Ar./Tt. Ar.<br>(%) | 43.21 +/- 1.64 | 42.90 +/- 1.72 | 41.12 +/- 1.80 \$* | 40.80 +/- 1.22 \$* |
| Ct. TMD<br>(mgHA/cm<sup>3</sup>) | 1208 +/- 8 | 1199 +/- 13 | 1197 +/- 12 | 1196 +/- 6 |
| Ct. Porosity (%) | 1.08 +/- 0.28 | 0.78 +/- 0.21 ^+ | 1.45 +/- 0.42 \$* | 1.21 +/- 0.34 +\$* |

The relatively mild effects of prednisolone on bone mass are corroborated by its small effects on serum markers of bone turnover measured at study completion (4 weeks after prednisolone pellet implantation). Prednisolone did not cause significant changes in serum procollagen type 1 N-terminal peptide (P1NP) or C-terminal collagen crosslinks (CTX) in AAV8-control or AAV8-Ostn-treated mice (Fig 3F, G). Taken together, these micro-CT and serum bone turnover marker data demonstrate only mild changes in cortical and trabecular bone mass and bone remodeling in response to prednisolone administration in this study. Mild, beneficial changes in cortical thickness were noted in prednisolone-treated mice that also received AAV8-Ostn.

Consistent with previous reports (33), we found strong mRNA expression of osteocrin in the tibial periosteum via *in situ* hybridization (Figure 4A, top), predominantly on the antero-medial bone surface. As in our humerus RT-qPCR data, a modest reduction in periosteal Ostn expression was noted in prednisolone-treated bones; fewer bones showed strong mRNA signal and no Ostn mRNA was detected in one prednisolone-treated mouse (Figure 4A, bottom). This suggests that normal Ostn signaling from periosteal cells to osteocytes is disrupted during glucocorticoid excess.

**Figure 4:**
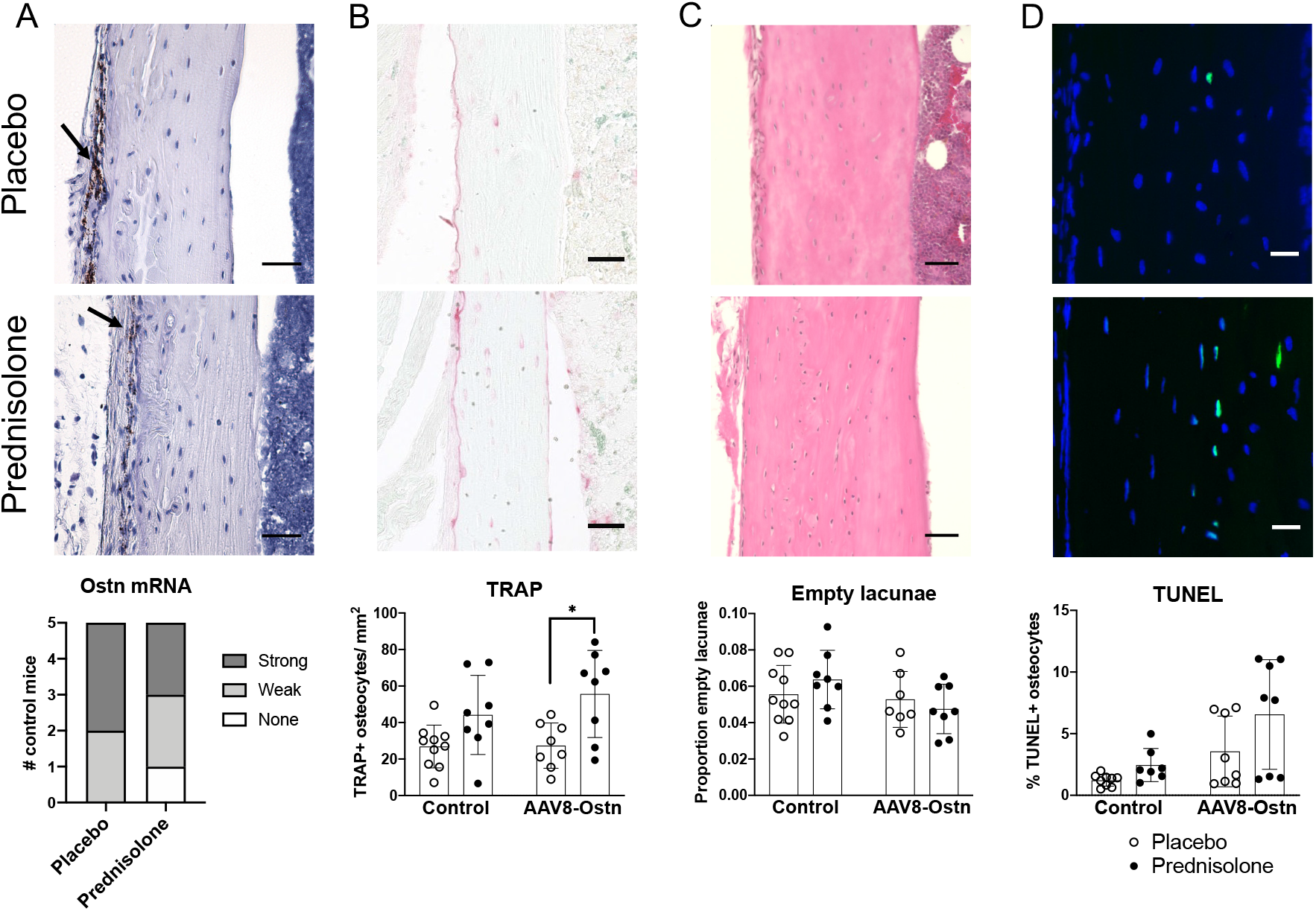
Prednisolone and AAV8-Ostn induce osteocyte apoptosis and TRAP expression in cortical bone. (A) Osteocrin mRNA expression was measured by *in situ* hybridization in AAV8-Control-treated cortical bone. Signal was semi-quantitatively analyzed but not statistically compared. Arrows indicate signal in periosteum. Scale bar = 50 μm. (B) Tartrate-resistant acid phosphatase (TRAP) expression was quantified as number of positively stained osteocytes per cortical bone area. Images show AAV8-Control groups. Scale bar = 50 μm. (C) Proportion of empty osteocyte lacunae was counted in hematoxylin and eosin-stained cortical bone. Images show AAV8-Control groups. Scale bar = 50 μm. (D) Terminal deoxynucleotidyl transferase dUTP nick end labeling (TUNEL) was used to quantify apoptotic osteocytes in cortical bone (one outlier removed from placebo+AAV8-Control group and one outlier removed from prednisolone+AAV8-Control group). Images show AAV8-Control groups. Scale bar = 20 μm. *p<0.05 compared to corresponding placebo group. All error bars show +/- SD.

Since endogenous expression of Ostn and the protective effects of AAV8-Ostn appeared to be limited to cortical bone, we conducted further histological analyses on cortical bone in the tibia. Consistent with RT-qPCR data in humeri, the number of TRAP-positive osteocytes was increased by prednisolone, though this difference only reached statistical significance in AAV8-Ostn mice (Figure 4B). Prednisolone did not significantly increase the frequency of empty osteocyte lacunae (Figure 4C) or the percentage of apoptotic (TUNEL-positive) osteocytes (Figure 4D), although TUNEL staining indicated trends towards increased apoptosis in response to both prednisolone and AAV8-Ostn. While the effects of prednisolone on osteocyte TRAP expression and viability were mild, they were not prevented by AAV8-Ostn.

GC excess leads to early, deleterious changes in osteocyte morphology (10,18,19). To visualize the effects of prednisolone and AAV8-Ostn on the osteocyte canalicular network, we used both silver nitrate (stains extracellular matrix and cellular structures) and phalloidin staining (stains filamentous actin). In AAV8-control-treated mice, prednisolone significantly reduced canalicular length in silver nitrate-stained bone (Fig 5A) and density of phalloidin-stained cell projections (Fig 5B). Neither of these metrics were significantly altered by prednisolone in AAV8-Ostn-treated mice, and canalicular number was not affected in either group. These results indicate a modest protective effect of AAV8-Ostn on osteocyte morphology that does not correlate with detectable changes in osteocyte apoptosis at this timepoint.

**Figure 5:**
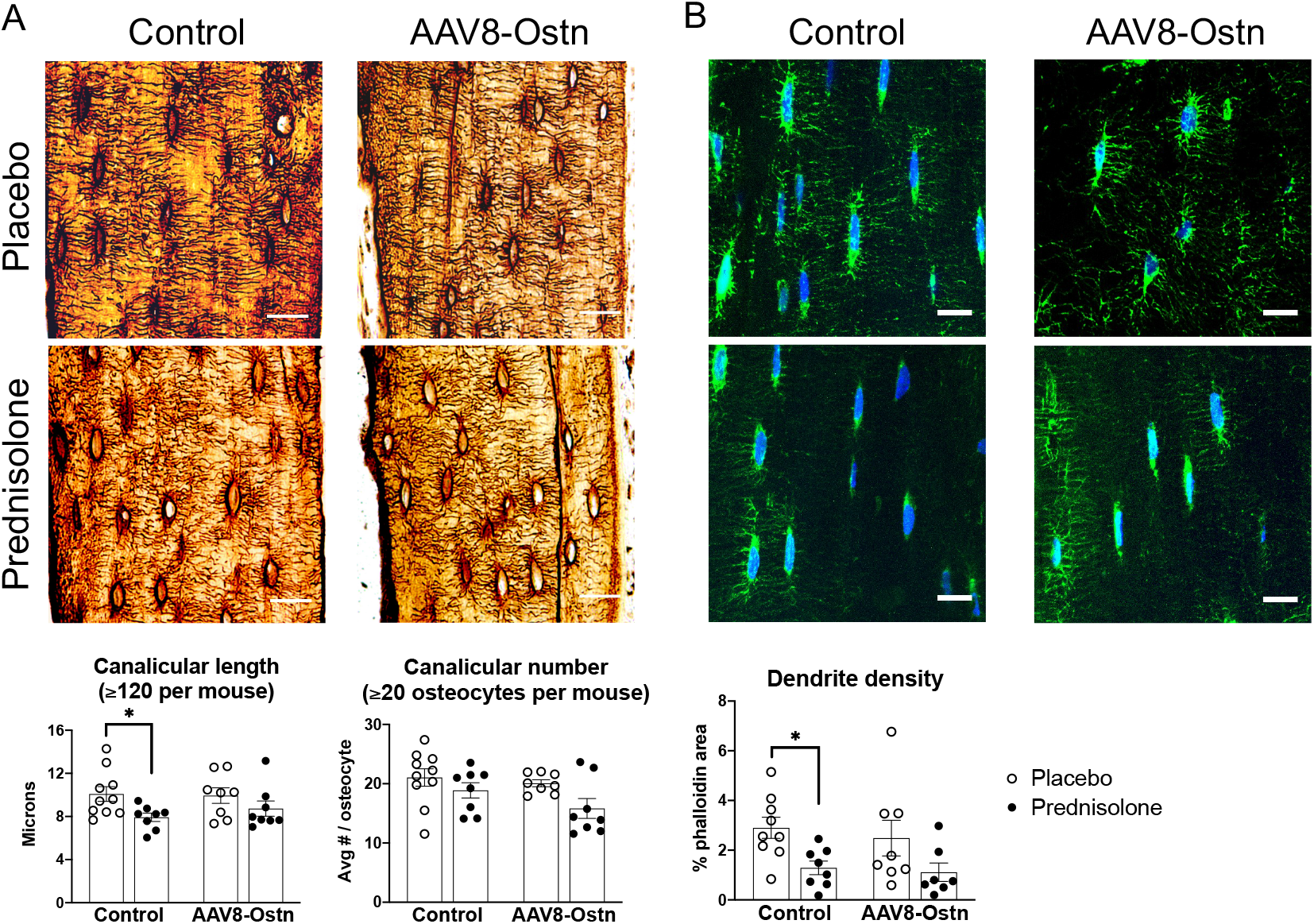
Prednisolone reduces canalicular length and dendrite density in AAV8-Control mice only. (A) Canalicular length and canalicular number were measured in silver nitrate-stained cortical bone. Scale bar = 20 μm. (B) Phalloidin staining of F-actin was used to measure density of osteocyte dendrites (one outlier removed from placebo+AAV8-Control group and one outlier removed from prednisolone+AAV8-Ostn group). Scale bar = 10 μm. *p<0.05 compared to corresponding placebo group. All error bars show +/- SEM.

## Discussion

This study tested the hypothesis that systemic treatment with liver-produced Ostn peptide could prevent the harmful prednisolone-induced changes to bone mass, osteocyte viability, and osteocyte morphology in cortical bone. After 4 weeks of prednisolone treatment, we found small changes in cortical thickness, canalicular length, and dendrite density that were significant only in control AAV8-treated mice. However, trabecular BMD and osteocyte TRAP staining were significantly changed by prednisolone in AAV8-Ostn treated mice, and cortical porosity and cortical area fraction were significantly changed by prednisolone irrespective of AAV8 treatment group. Therefore, the apparent protective effects of Ostn on osteocyte morphology did not translate to functional improvements in osteocyte maintenance of bone. While Ostn and other ligands of Npr2/Npr3 may be useful at different doses or in different skeletal diseases, this study does not support a dramatic therapeutic benefit of Ostn to combat deleterious skeletal effects of glucocorticoid excess.

Our hypothesis was supported by several studies showing beneficial effects of Ostn and CNP in bone and growth plates of developing mice. High circulating Ostn in young mice stimulates bone growth in a manner that depends on NPR-C and endogenous CNP expression (34). Likewise, enhancing CNP signaling through NPR-B regulates bone length, mineralization, and strength in growing mice (35). In mice lacking Sp7 expression in osteocytes, AAV8-Ostn treatment from 3-6 weeks old preserves osteocyte morphology, osteocyte viability, and cortical porosity (20). Recently, Watanabe-Takano et al. described that Ostn is involved in bone formation during recovery from unloading, suggesting an additional role for Ostn in mature bone (33).

Here we investigated whether mature osteocytes could also benefit from Ostn in a disease model. As in previous studies, this experiment used the liver to overexpress Ostn and deliver the peptide to bone (20,34). We found that AAV8 infection of the liver caused rapid upregulation of *Ostn* mRNA that was still apparent at 35 days post-injection. To confirm that Ostn successfully targeted and induced signaling in bone, we measured bone expression of *Fos*, which was proportional to hepatic *Ostn* expression. In contrast to treatment of young developing mice (3-6 weeks old) lacking Sp7 in osteoblasts and osteocytes (20), osteocrin treatment here (7-12 weeks old) had few effects on osteocytes or bone. We observed that, as expected, AAV8-Ostn reduced cortical porosity. However, because AAV8-Ostn reduced osteocyte apoptosis and improved osteocyte morphology defects in Sp7 conditional knockout mice, we were surprised to see that Ostn tended to increase TUNEL staining in osteocytes here. The protective effect of osteocrin on canalicular length was somewhat more striking in the anterior tibia, so it is possible that osteocrin exerts site-specific effects related to cortical drift during development or proximity to endogenous osteocrin expression. These age and site-dependent differences will be important to consider as the mechanism and function of osteocrin are studied further.

An important limitation of the current study is the mild overall effect of prednisolone on control mice. At similar reported doses of prednisolone and 3-6 weeks of treatment, other studies have reported more robust changes in trabecular BV/TV, serum bone turnover markers, and osteocyte viability than were detected here (13,14,16,18,30,36,37). Changes in mouse body weight throughout the study period give us confidence in the efficacy of the glucocorticoid excess model, and we did find the expected prednisolone-induced changes in cortical thickness, cortical porosity, and osteocyte morphology. Some of these outcomes may be time-dependent. For example, suppression of PLR genes was previously reported at 7 days and may be resolved or compensated for by 28 days (18), and osteocyte viability would perhaps be more significantly different after a longer treatment period (15). Therefore, future time course studies may be needed to assess effects of AAV8-Ostn on distinct prednisolone-induced skeletal changes. In addition, given the mild overall effect of prednisolone on our control mice, we cannot exclude the possibility that a higher dose/stronger overall effect of prednisolone would induce significant changes in mice treated with AAV8-Ostn. There were some apparent though statistically insignificant effects of prednisolone in AAV8-Ostn mice, and a larger effect in control mice or a larger sample size may tip the scales towards statistical significance in this group.

Glucocorticoid-induced osteoporosis is a major cause of skeletal fragility and fractures. GCs effectively induce a ‘perfect storm’ of changes in bone cells that culminates in increased bone resorption, reduced bone formation, and poor bone quality (1). Defects in osteocyte activity and morphology and subsequent osteocyte apoptosis occur early in the course of GC therapy and likely play a central role in driving subsequent changes in remodeling on bone surfaces. Therefore, treatments that restore osteocyte morphology and viability represent a promising novel approach to meet this important unmet medical need. Here we showed that systemically-administered Ostn (via AAV8 gene therapy) could improve cortical thickness and osteocyte morphology, but other GC-induced changes were not rescued by AAV8-Ostn in this study. It remains possible that different molecular pathways lead to osteocyte dysfunction in the settings of Sp7 and GC treatment. Despite the aforementioned limitations, these findings demonstrate the therapeutic potential for AAV8-Ostn for common diseases caused by osteocyte dysfunction and support further study into the translational potential of different modalities of bone-directed Ostn therapy.

## Acknowledgements

We thank all members of the Wein laboratory for stimulating discussions. MNW acknowledges funding support from the MGH Department of Medicine (Transformative Scholars award), the American Society of Bone and Mineral Research (Rising Star Award), and the National Institute of Health (AR067285). CMM acknowledges support from the NIH (T32DK007028). M.N.W. and CMM acknowledge generous support from Louise Pearl Corman, Ph.D. Micro-CT and bone histology preparation were conducted by the Center for Skeletal Research, an NIH-funded program (P30 AR075042). Confocal microscopy was supported by the NIH Shared Instrumentation Grant (SIG) S10OD021577.

---

**Supplemental Figure 1:**
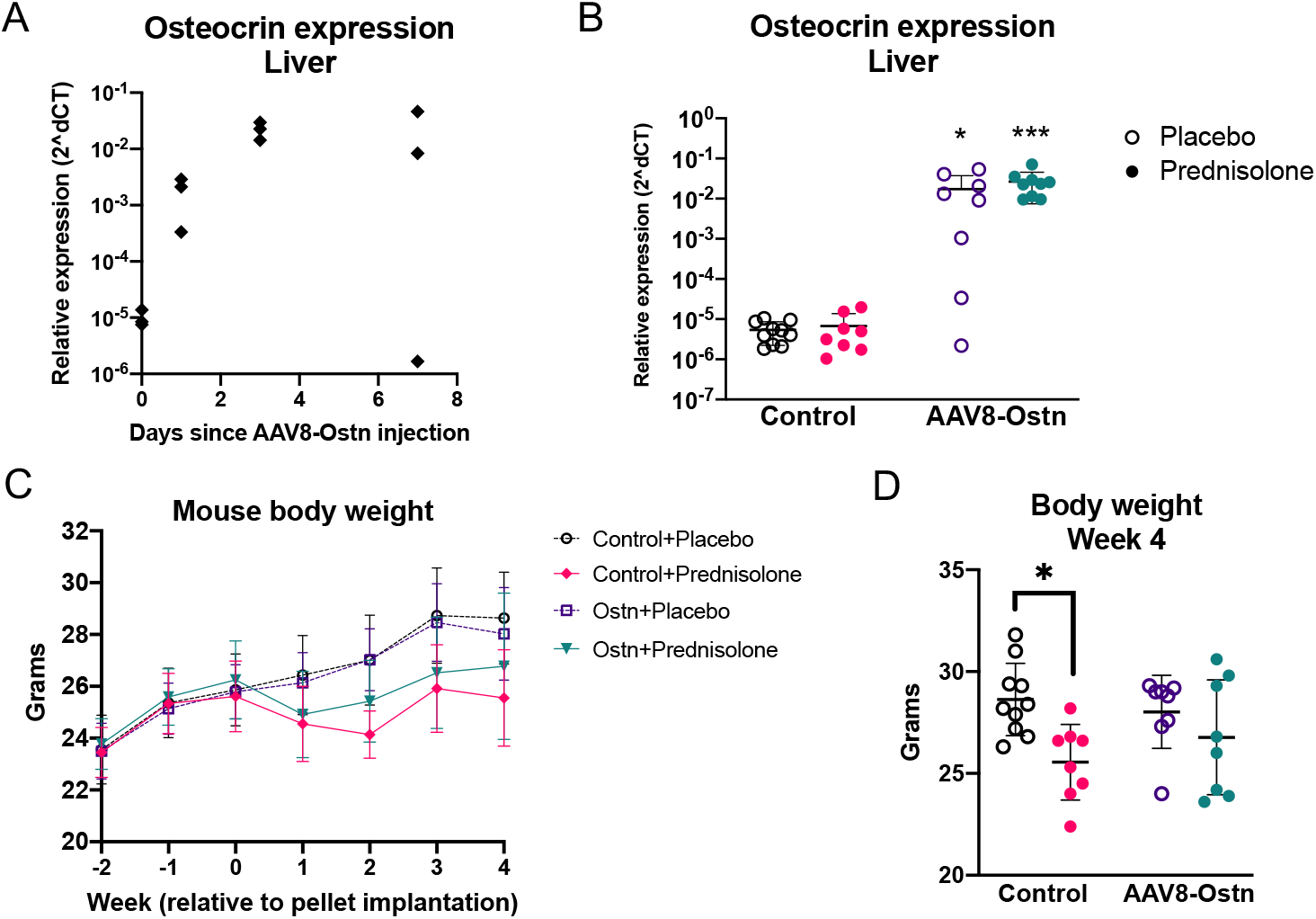
Systemic effects of AAV8-Ostn and prednisolone in mice. (A) Hepatic expression of *Ostn* mRNA was measured relative to *Actb* 1, 3, and 7 days after intraperitoneal injection of AAV8-Ostn in C57BL/6J mice. (B) Hepatic expression of *Ostn* mRNA was measured relative to *Actb* 35 days after intraperitoneal injection of AAV8-Ostn or AAV8-Control and 28 days after placebo or prednisolone pellet implantation in FVB mice. *p<0.05, ***p<0.001 relative to corresponding AAV8-control group. (C-D) Mice were weighed weekly starting 2 weeks prior to GC pellet implantation, and statistical comparisons were performed at the end of the experiment. *p<0.05 compared to corresponding placebo group. All error bars show +/- SD.

**Supplemental Figure 2:**
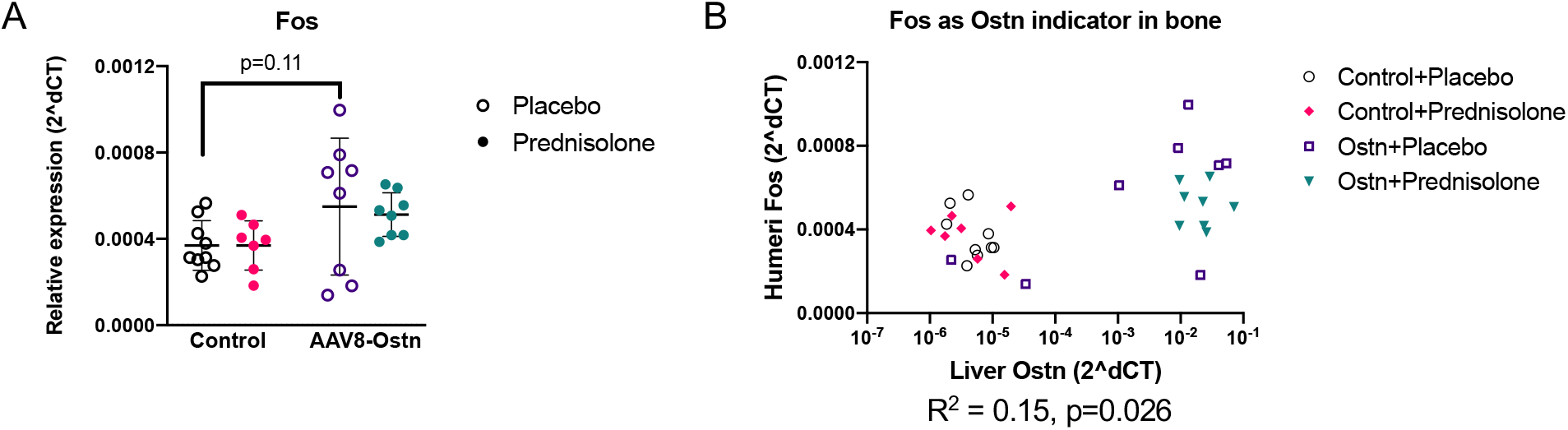
Osteocrin-induced changes in gene expression. (A) Expression of Erk1/2-responsive gene *Fos* was measured in humeri and normalized to *Actb* 35 days after intraperitoneal injection of AAV8-Ostn or AAV8-Control. Error bars show +/- SD. (B) *Fos* expression in humeri (normalized to *Actb*) is plotted relative to hepatic *Ostn* expression (normalized to *Actb*) for each mouse (R^2^ = 0.15, p=0.026).

